# Strain concentration drives the anatomical distribution of injury in acute and chronic traumatic brain injury

**DOI:** 10.1101/2024.05.22.595352

**Authors:** Adnan A. Hirad, Doran Mix, Arun Venkataraman, Steven P. Meyers, Bradford Z. Mahon

## Abstract

Brain tissue injury caused by mild traumatic brain injury (mTBI) disproportionately concentrates in the midbrain, cerebellum, mesial temporal lobe, and the interface between cortex and white matter at sulcal depths ^1–12^. The bio-mechanical principles that explain why physical impacts to different parts of the skull translate to common foci of injury concentrated in specific brain structures are unknown. A general and longstanding idea, which has not to date been directly tested in humans, is that different brain regions are differentially susceptible to strain loading^11,13–15^. We use Magnetic Resonance Elastography (MRE) in healthy participants to develop whole-brain bio-mechanical vulnerability maps that independently define which regions of the brain exhibit disproportionate strain concentration. We then validate those vulnerability maps in a prospective cohort of mTBI patients, using diffusion MRI data collected at three cross-sectional timepoints after injury: acute, sub-acute, chronic. We show that regions that exhibit high strain, measured with MRE, are also the sites of greatest injury, as measured with diffusion MR in mTBI patients. This was the case in acute, subacute, and chronic subgroups of the mTBI cohort. Follow-on analyses decomposed the biomechanical cause of increased strain by showing it is caused jointly by disproportionately higher levels of energy arriving to ‘high-strain’ structures, as well as the inability of ‘high strain’ structures to effectively disperse that energy. These findings establish a causal mechanism that explains the anatomy of injury in mTBI based on *in vivo* rheological properties of the human brain.

## Introduction

Mild traumatic brain injury (mTBI), or concussion, affects up to 3.8 million Americans annually^16^. In the acute phase, mTBI is associated with immediate symptoms, such as dizziness and headaches, and signs, such as loss of consciousness, memory loss, and oculomotor dysfunction. Prior research shows that the pattern of tissue disruption in TBI of all severities, including in mTBI, is to some degree stereotyped: regardless of where the skull is impacted, tissue injury is observed to disproportionaterly involve the midbrain, cerebellum, mesial temporal lobe structures, as well as the interface between cortex and white matter at sulcal depths ^1–12^. Those midline structures tend to underpin functions related common signs and symptoms of mTBI, including consciousness, balance, oculomotor function, and memory ^11,12^. Tissue disruption can result in a cascade of electrophysiological and neurochemical dysfunction, and thus the physiological basis of the clinical signs and symptoms of m/TBI is complex and multifactorial^17^. Nonetheless, the original cause of dysfunction and long-term pathology can be traced to how a physical impact to the head translates to a *stereotyped* pattern of brain injury in the living human brain ^11–13,15,17–25^. Here we test whether that stereotyped injury pattern is predicted by independent measures of which brain structures exhibit high strain, or shear wave concentration, under skull loading. Advances in understanding how shear waves to the skull cause energy to concentrate in specific structures of the brain are crucial for developing a mechanistic model of TBI injury. A biomechanical model of how m/TBI causes brain tissue injury has broad implications for advancing basic biological understanding, and represents a critical step for developing preventative, therapeutic, and prognostic tools.

In the context of mTBI, tissue disruption stems from the deformation, specifically shear strain, encountered by brain tissue in response to a head impact ^14,15,26^. Strain (*ɛ*
) is a quantifiable mechanical response of a structure when subjected to a load (in this paper we are only concerned with shear strain, and use strain and shear strain interchangeably). Relatively high strain (strain concentration) in specific brain structures could be due to elevated levels of shear wave forces, signifying a differentially high level of energy transfer to those structures. A non-exclusive alternative is that regions stereotypically associated with injury in m/TBI exhibit differential stiffness or damping compared to the surrounding brain structures, such that, for a given level of force loading, those brain structures deform more easily. These possibilities can be independently evaluated by taking advantage of basic reheological principles, as applied to in vivo Magnetic Resonance Imaging data in humans.

Energy transferred to a viscoelastic medium, such as the brain, can be deformative, or it can be dissipated as heat. Shear strain energy is proportional to the product of the shear stiffness and shear strain of that region. Storage and loss moduli are measures that characterize the deformative and dissipative behaviors of a viscoelastic structure. Both the storage and loss moduli of the brain increase as the frequency of applied forces increases. The relative rate of deformation and energy dissipation of brain tissues can thus be captured by the slopes of the storage and loss moduli as a function of impact frequency. That slope (dispersion) serves as an index of the degree to which the region in question can dissipate the energy that arrives to it. Dispersion, which is driven by local structural and material properties^27^, thus holds real promise as a quantitative index about how a particular brain region handles the impact energy transferred to it. Consequently, regions where greater impact energy leads to relatively higher strain despite high stiffness are likely to sustain tissue disruption compared to regions with similar strain but lower stiffness, or similar stiffness, but lower strain. As such, analysis of strain energy holds tremendous potential for adjudicating the factors contributing to a stereotyped pattern of tissue injury in m/TBI.

In summary, if shear strain is indeed the primary factor driving injury in mTBI, then the pattern of tissue disruption in mTBI should mirror the distribution of shear strain concentration within the brain. In this paper we test three specific empirical predictions: 1) A whole-brain strain concentration map will identify brain regions known in the literature to exhibit stereotyped vulnerability to concussive impacts, including the midbrain, cerebellum, mesial temporal lobe structures, and sulcal depths; 2) strain concentration in midline regions will be independent of the location of mechanical stimulation of the skull (comparing anterior-posterior vs. lateral-medial directions of stimulation); and 3) to demonstrate construct validity that strain concentration measured with MRE provides a vulnerability map relevant to overt mTBI injury, we test how strain concetration predicts tissue disruption in separate cohorts of mTBI patients in the acute, subacute and chronic phases after injury. We further seek evidence for each of the two non-exclusive potential causes of high strain concentration: whether high strain concentration derives from the differential focusing of shear waves in high strain regions, and/or the inability of those regions to effectively dissipate the energy that arrives. To test these predictions, we used Magnetic Resonance Elastography (MRE) coupled with non-injurious perturbation of the skull in healthy control participants to generate a population-level whole-brain strain concentration map. That strain concentration map was then used to predict the location of tissue disruption in a new cohort of patients with clinically confirmed mTBI. Tissue disruption was quantified in mTBI participants using Apparent Fiber Density (AFD), a diffusion MRI-based metric that indexes tissue integrity in both white and gray matter^28, 29^.

## Methods

### Subacute and chronic mTBI Participants

We conducted retrospective analysis of high school and college athletes with a history of concussion and Post Concussive Syndrome (PCS) who had presented between February 2016 and December 2019. All subjects in our study were referred to the University of Rochester Medical Center outpatient imaging center due to incomplete resolution of symptoms following concussion. All subacute and chronic mTBI patietns and controls were scanned on the same 3T Skyra MRI scanner. Inclusion criteria consisted of a history of concussion and PCS, while exclusion criteria included dental braces, prior brain surgery, ventricular shunt, skull fractures, or other standard contraindications for MR imaging. Diagnosis of concussion was made by ED physicians initially (recovered from the medical record). PCS was determined through standard clinical consensus across a multi-disciplinary team of neurologists, physical medicine and rehabilitation physicians, and sports medicine physicians. All diagnoses (initial mTBI and subsequent PCS) were made independent of, and blinded to, hypothesis formulation and subsequent inclusion in the analyses of the current report. The control group consisted of age-matched participants with no history of concussion. The University of Rochester Medical Center Institutional Review Board approved the retrospective analysis of structural (T1) and dMRI data for all subjects presenting with concussion from 2012 to 2022. The data were accessed by a single clinician (SM), and from that point, it was de-identified. The data curation was conducted between June and December of 2022. None of the other study members had access to the identities of the subjects at any point. Additionally, SM had no access after the initial identification of subjects. Number of previous concussions, loss of consciousness at most recent concussion, and time between injury and MRI were determined from review of the electronic medical records. An experienced neuroradiologist (SM) reviewed all MRI examinations, blinded to subsequent hypothesis testing, for any artifacts that might affect study quality, and for the presence of intracranial hemorrhage, signal abnormalities in the brain, hydrocephalus, and congenital or developmental anomalies. That MRI review was performed blinded to all analyses and findings using MRE to identify regions of differneitally high versus low strain. 54 particpants had dMRI data as well as known time between injury and imaging and could therefore be classified as subacute (delay between injury and imaging < 90 days) and chronic (delay > 90 days) cohorts. 23 subjects (age (mean ± SD) – 20.4±2.7, 9 males, 14 females) were in the subacute time period at the time of dMRI data acquisition, and 31 were in the chronic time period (age (mean ± SD) – 20.7±3.8, 16 males, 15 females). The University of Rochester Institutional Review Board approved this study, and written informed consent was obtained from all participants.

### Acute mTBI Cohort

We conducted retrospective analyses on 15 male and 14 female mTBI patients (mean age = 19.5, median age = 19) and 15 male controls (mean age = 21.6, median age = 21). The individuals diagnosed with concussion represented a subset of a broader group of NCAA contact-sport athletes at the University of Rochester and the Rochester Institute of Technology who were monitored for concussion. Between 2009 and 2014, National Collegiate Athletic Association (NCAA) division I and III collegiate contact sport athletes were followed prospectively for a diagnosis of mTBI ^6,58,59^. mTBI was defined as an injury witnessed by an on-field certified athletic trainer and meeting the definition of concussion as defined by the Sport Concussion Assessment Tool 2. The diffusion MRI scans were collected within 72 hours post-injury on concussed individuals^6^. As such, diagnosis of mTBI was made independent of decisions about including those participants in the current study. The University of Rochester Institutional Review Board approved this study, *as an exempt study for secondary use of pre-existing data*.

### MRI Acquisition for subacute and chronic mTBI cohorts and matched controls

MR imaging was conducted on a Siemens Skyra 3T scanner using a 20-channel head and neck coil. Diffusion imaging was performed with a b-value of 1000 s/mm^2^, using 64 diffusion-encoding directions. In addition, a b=0 s/mm^2^ image was collected. Additional dMRI parameters included: FOV = 256 x 256 mm, 70 slices, image resolution = 2×2×2 mm^3^, TR/TE = 9000/88 ms, GRAPPA factor = 2, acquisition time = 10 minutes and 14 seconds. A GRE sequence was also collected with TEs = 4.92, 7.38 ms at the same resolution of the dMRI to correct for susceptibility-induced distortion. The MRI protocol also consisted of T1 magnetization-prepared rapid gradient echo (MP-RAGE) (TR = 1200 ms, TE= 2.29 ms, TI = 600 ms, flip angle =8 degrees, FOV = 250 mm, 1 x 1 x 1 mm^3^, 208 slices), 3D Axial SWI (TR = 27 ms, TE = 29 ms, FOV = 220 mm, 1.5 mm slice thickness, 88 slices), and Double IR fat suppressed FLAIR (TR = 7500 ms, TE = 321 ms, TIs = 3000, 450 ms, FOV = 260 mm,1.4 mm slice thickness, 120 slices), which were not used in the current study, but were collected as part of standard clinical care.

### General Procedures and MRI Acquisition Parameters for the Acute mTBI Cohort

Participants were tested on a Siemens 3T Tim Trio scanner using a 32-channel head coil at the Rochester Center for Brain Imaging (now named: Center for Advanced Brain Imaging and Neurophysiology). High-resolution structural T1 contrast images were acquired using a magnetization-prepared rapid gradient echo (MP-RAGE) pulse sequence at the start of each participant’s first scanning session (TR = 2530 ms, TE = 3.44 ms, flip angle = 7 degrees, FOV = 256 mm, matrix = 256 x 256, 1×1×1 mm sagittal left-to-right slices). DTI sequence parameters were: TR/TE =10 s/89 ms, voxel size 2×2×2 mm, 60 diffusion directions with b=1000 s/mm2 and 10 averages of b=0.

### Diffusion MR Analysis

Fieldmaps were generated from the GRE data using the fugue command in FSL^70^. The fieldmaps were unwrapped and used to correct susceptibility-induced distortions. The FSL eddy command was used to correct for effects of eddy currents as well as subject motion^71^. The data was subsequently bias-field corrected using ANTs (http://stnava.github.io/ANTs/). *dipy* toolbox version 1.1.1 (https://www.dipy.org) was used to calculate DTI metrics and apparent fiber density (AFD). AFD a marker that is related to radial diffusivity, and is more sensitive in regions with multiple fiber orientations (see Raffelt et al^29^), depends on axon diameter and packing density^29^ and resolves crossing fibers at b=1000 ^28^. For detailed review of AFD calculation see Raffelt and colleagues ^29^.

### MRE Data Acquisition

MRE data were obtained from publically available depositions collected in two separate cohorts of healthy control participants. The first cohort consisted of 59 subjects and the data were collected at, and made publicly available by, the University of Delaware (UDEL) using a Siemens 3T Prisma scanner with a 64-channel head/neck coil (complete protocol available at ref^30^). Skull displacements were generated using a resoundant acoustic (Resoundant Acoustic Driver System, Resoundant™ Rochester, MN) driver system with a soft pillow occipital actuator. Three separate scans were acquired with 30, 50 and 70 Hz occipital skull actuation. A 3D multiband, multishot spiral sequence was used to measure brain tissue displacements, providing whole-brain coverage with an imaging resolution of 1.5 mm isotropic (240 x 240 x 120 mm³).

To estimate the complex shear, storage, and loss moduli, an NLI inversion algorithm was utilized ^72^. Shear stiffness was calculated from those properties. Additionally, octahedral shear strain (OSS) was derived at each voxel, representing the maximum shear strain regardless of direction ^73^. Shear strain energy was calculated as the product of stiffness and OSS at the voxel level 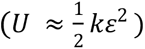, where U = energy, k = shear stiffness, and *ɛ* = *strain*.

The second MRE cohort consisted of 24 subjects, and the MRE data were obtained from the same public repository, but collected at the University of Washington St. Louis (WUSTL) ^30^. Participants underwent harmonic lateral skull actuation at 50Hz using the same Resoundant actuator system using a passive flexible silicone bottle at the lateral skull. The lateral actuation approach provided a different pattern of skull motion compared to the occipital actuation used in the UDEL dataset. MRE data for the WUSTL cohort was acquired at a slightly lower isotropic voxel resolution of 3 mm, compared to the 1.5 mm isotropic resolution of the UDEL cohort.

### MRE Processing and analysis

The analysis was conducted in MNI (Montreal Neurological Institute) space to facilitate the generation of group-level strain concentration maps and support extensionality for testing in the separate cohorts of mTBI participants. Each participant’s MPRAGE and MRE magnitude images underwent brain extraction using the BET (Brain Extraction Tool) of FSL (FMRIB Software Library). MPRAGE images were transformed and warped into MNI space using the ANTs (Advanced Normalization Tools, http://stnava.github.io/ANTs/) SyN tool. Similarly, each subject’s MRE magnitude image was transformed and warped into their native MPRAGE space. The resulting transformation matrices from the previous steps were concatenated and applied to the MRE data, bringing those data into MNI space. This transformation allowed for the analysis of strain concentration at the group level, enabling comparisons and statistical evaluation across participants and cohorts.

## Results

### Rheological analyses identify brain regions with high strain concentration

MRE data were analyzed from 59 healthy control subjects who underwent occipital mechanical stimulation at 50 Hz, and 24 subjects who underwent lateral mechanical stimulation at 50 Hz^30^. Our core analyses were focused on 50Hz occipital stimulation, because it is the most commonly used frequency for MRE data collection, and it corresponds to the largest possible sample size involving both occipital and lateral stimulation. The dependent variable, Octahedral Shear Strain (OSS), was normalized across all voxels within each subject prior to group statistical analysis in stereotactic space. Separate group-level analyses were conducted for the lateral and occipital actuation data, consisting of voxel-wise t-maps (two-tailed, FDR corrected at q < .05). For hypothesis testing, regions of differentially high strain concentration (High Strain Regions, HS) were defined as voxels at or above t = 4.0 (FDR-corrected q < .05). We selected a t-score threshold of 4 as it survives False Discovery Rate (FDR) correction. Voxels with levels of strain below that threshold were defined as low strain regions (LS). For both lateral and occipital mechanical stimulation, the midbrain, cerebellum, and mesial temporal lobe (hippocampus and parahippocampus) structures exhibited differentially high strain concentration (Figure 1A and B). Furthermore, and in agreement with prior histopathologic findings on post-mortem resection in chronic traumatic encephalopathy (CTE) ^12,31^, the depths of the sulci as opposed to gyral crowns, also exhibited differentially high strain.

**Figure 1:**
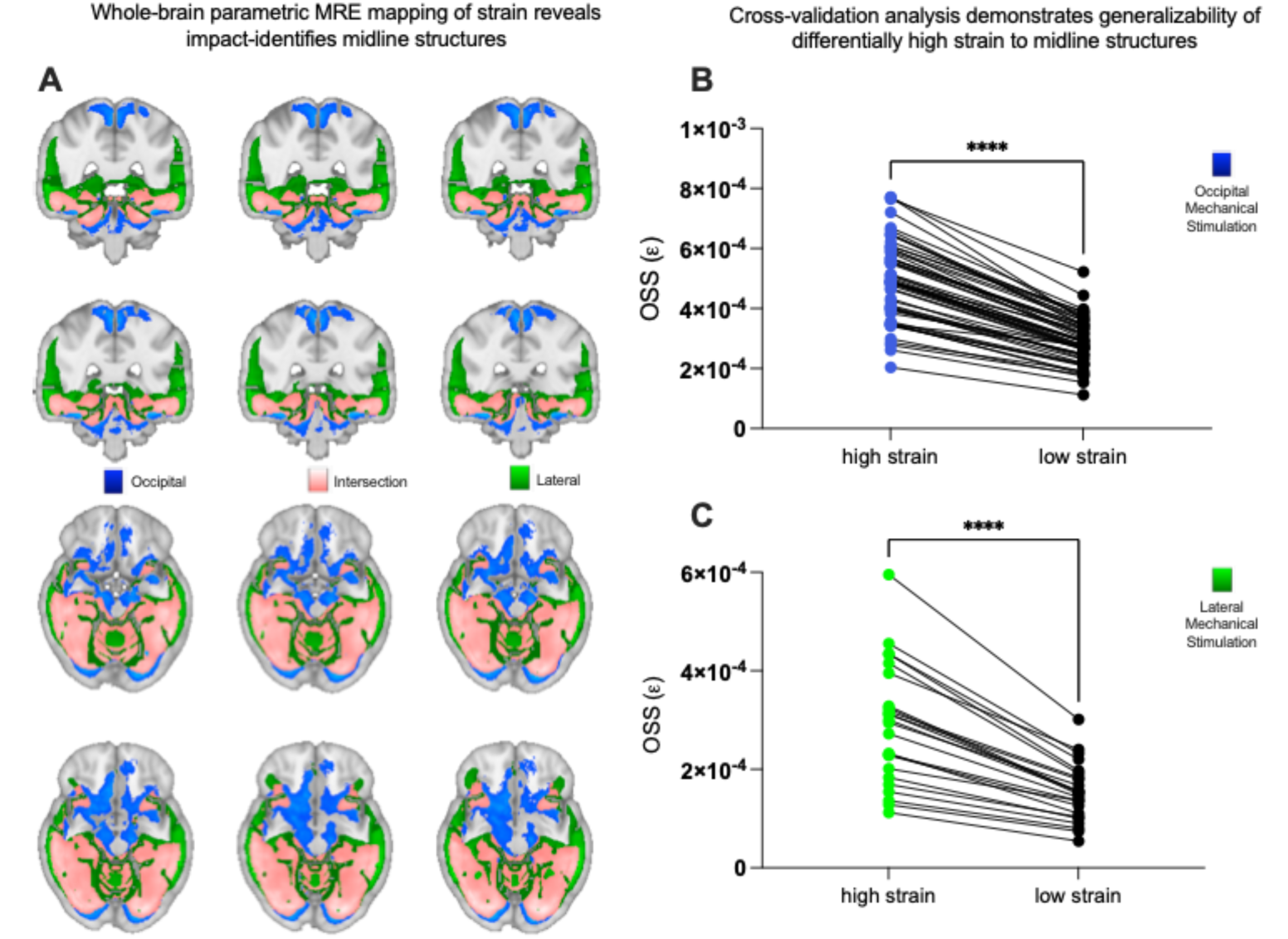
Differentially high strain concentration in specific brain structures caused by 50Hz occipital and lateral actuation of the head, measured using MRE. ***A)*** Strain concentration from occipital head actuation (blue), lateral head actuation (green), and regions of intersection (pink) between occipital and lateral actuation. Pink regions experience differentially high strain concentration that is invariant to the location of skull actuation. ***B)*** Leave-one-out cross-validation analysis of the occipital actuation MRE data (n=59) demonstrating differentially increased strain concentration across *all* subjects. ***C)*** Leave-one-out cross-validation analysis of the lateral actuation MRE data (n=24) demonstrating strain concentration across *all* subjects.

### Cross-validation analysis

In order to quantify the generalizability of the Strain Concentration Analysis, we conducted a leave-one-out cross-validation of the lateral and occipital mechanical stimulation data. High Strain Regions were defined at t = 4.0 (FDR-corrected q < .05) on group data from n-1 participants, and the resulting ROIs for high and low-strain regions were used to extract OSS from the left-out participant (folded over all participants). The results indicate a robust and consistent pattern of differentially high strain concentration in the respective regions, across all subjects, for both lateral (t = 10.3, p < .0001) and occipital actuation (t = 23.5, p < 0.0001; Wilcoxon matched-pairs rank test, Fig 1 B and C). These data support the existence of stereotyped patterns of high strain concentration in specific brain structures that is independent of whether the mechanical stimulation of the skull was lateral or occipital.

### High strain concentration regions predict differential tissue injury in acute, subacute, and chronic mTBI

To test whether regions that exhibit differentially high strain concentration also exhibit injury in mTBI, we analyzed diffusion MRI (dMRI) data from mTBI patients scanned within 72 hours of injury (acute, n = 29), between 2 weeks and 90 days after injury (subacute, n = 31), and more than 90 days post-injury (chronic, n = 23). For each mTBI cohort, dMRI data from the same scanner, and scanning parameters, were collected from an age-matched control group without a history of TBI. We extracted Apparent Fiber Density (AFD) at each voxel in each subject (mTBI participants and matched controls). AFD values were averaged for regions identified in the MRE analyses as exhibiting high strain and separately for regions exhibiting low strain. High Strain Concentration regions were defined based on MRE data as the intersection of high-strain regions observed *both for* posterior (occipital) and lateral skull perturbations. Given the inherent variability across participants in diffusion MRI metrics like AFD, each mTBI and control subject’s dMRI values from high-strain regions were normalized to the respective AFD from low-strain regions. This analysis has high rigor and reproducibility because ROIs are defined in a group of 83 healthy controls using stringent criteria and then tested (in stereotactic space) in three independent TBI cohorts, covering the full timeline of post-concussive recovery (acute through chronic). Three planned 2_[high vs. low strain]_ X 2_[mTBI vs control]_ ANOVAs, with AFD as the dependent measure, were conducted for the acute, sub-acute, and chronic mTBI patients. Critically, the interaction was significant for all three post-concussive groups: acute (F = 4.63; P < 0.037), subacute (F = 60.79; P < 0.0001), and chronic (F = 31.38; P < 0.0001). We then tested the directed hypothesis that the cause of the significant interactions was differentially lower AFD in mTBI participants in regions of high strain concentration compared to regions of low strain concentration, compared to the same contrast in controls: [mTBI_[high strain ROI]_ > mTBI_[low strain ROI]_] > Control_[high strain ROI]_ > Control_[low strain ROI]_]. That directed hypothesis test was significant for each of the mTBI groups acute vs. control (t = 3.0; p < 0.005) Fig 2A; subacute vs control (t = 9.0, p < 0.0001) and chronic vs control (t = 6.6; p < 0.0001) Fig 2B).

**Figure 2:**
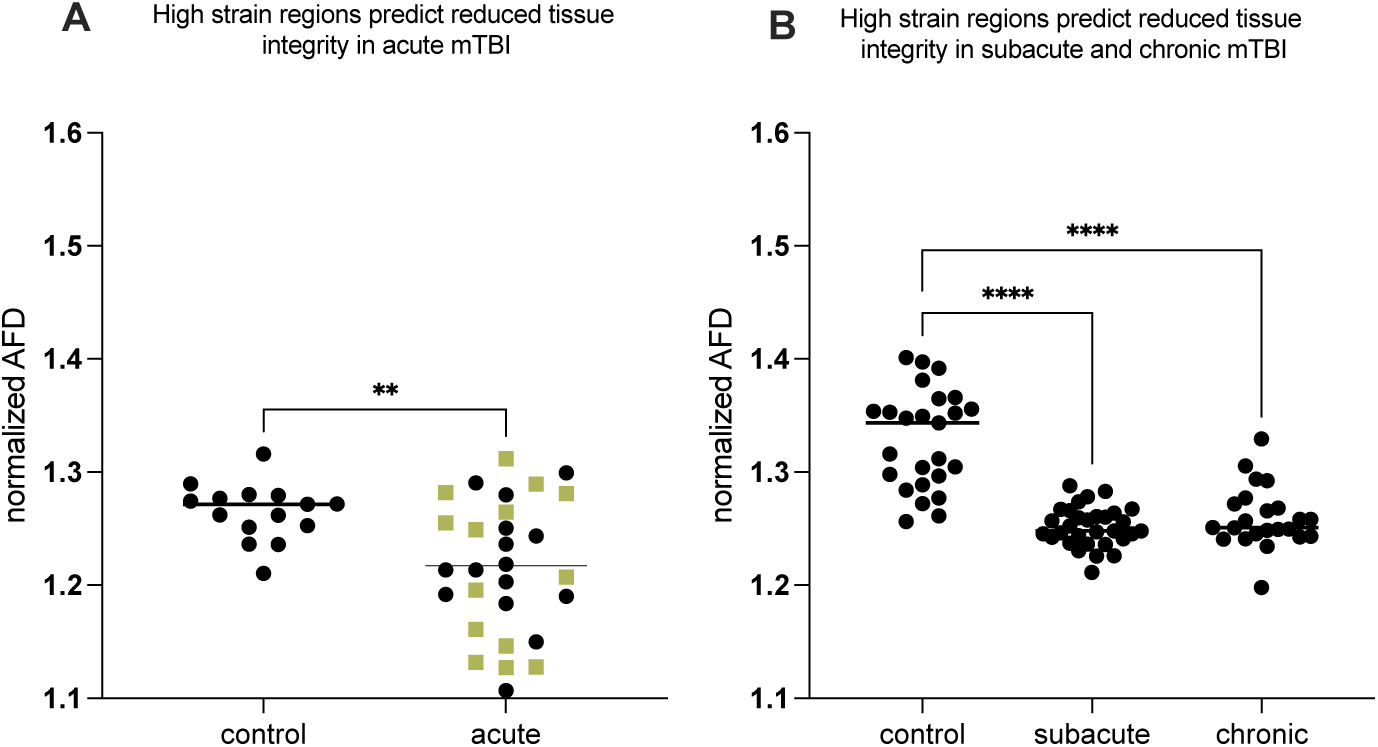
High strain concentration, measured in healthy participants using MRE, predicts regions of reduced tissue integrity in mTBI patients. Y axes represent AFD values in high strain regions normalized to low strain regions (within particpants). As described in the text, 2 (mTBI vs Controls) x 2 (High- vs Low-Strain Regions) ANOVAs demonstrated significant interactions in each mTBI sub-group. *A)* There was significantly lower Apparent Fiber Density (AFD) in high-strain regions normalized to low-strain regions (AFD_HS_/AFD_LS_) in acute mTBI compared to control subjects (olive green squares are female mTBI patients, separated because the controls are entirely male). *B)* There was differentially lower AFD in high-strain regions normalized to low-strain regions (AFD_HS_/AFD_LS_) in subacute and chronic mTBI participants, compared to control subjects.

We then sought to examine the relation between strain concentration and changes in structural integrity, broadly and without the imposition of a statistical threshold on the whole-brain map to separate high versus low strain regions. We assessed the relation, across all voxels in the brain, between strain concentration and AFD. Specifically, for the MRE data, Octahedral Shear Strain (OSS) values were normalized across all voxels within each participant. For the mTBI diffusion data, at each voxel the average AFD from control participants was subtracted from the corresponding average AFD from the mTBI participants—resulting in a measure of differential reduction of AFD in mTBI compared to controls. Importantly, this approach normalizes for the expected high variability of AFD across voxels within a brain. OSS values (averaged across all participants) for all voxels were binned into 0.25 Z-Score intervals spanning the entire range of values in the OSS data. Those bins were used to compute the median differences in AFD between mTBI and controls. The analysis demonstrated, at the whole brain level, that brain regions/voxels with higher strain also exhibit greater reduction in tissue integrity in mTBI patients compared to controls (Fig 3). This threshold-agnostic approach reinforces the relation between strain concentration and injury in mTBI.

**Figure 3:**
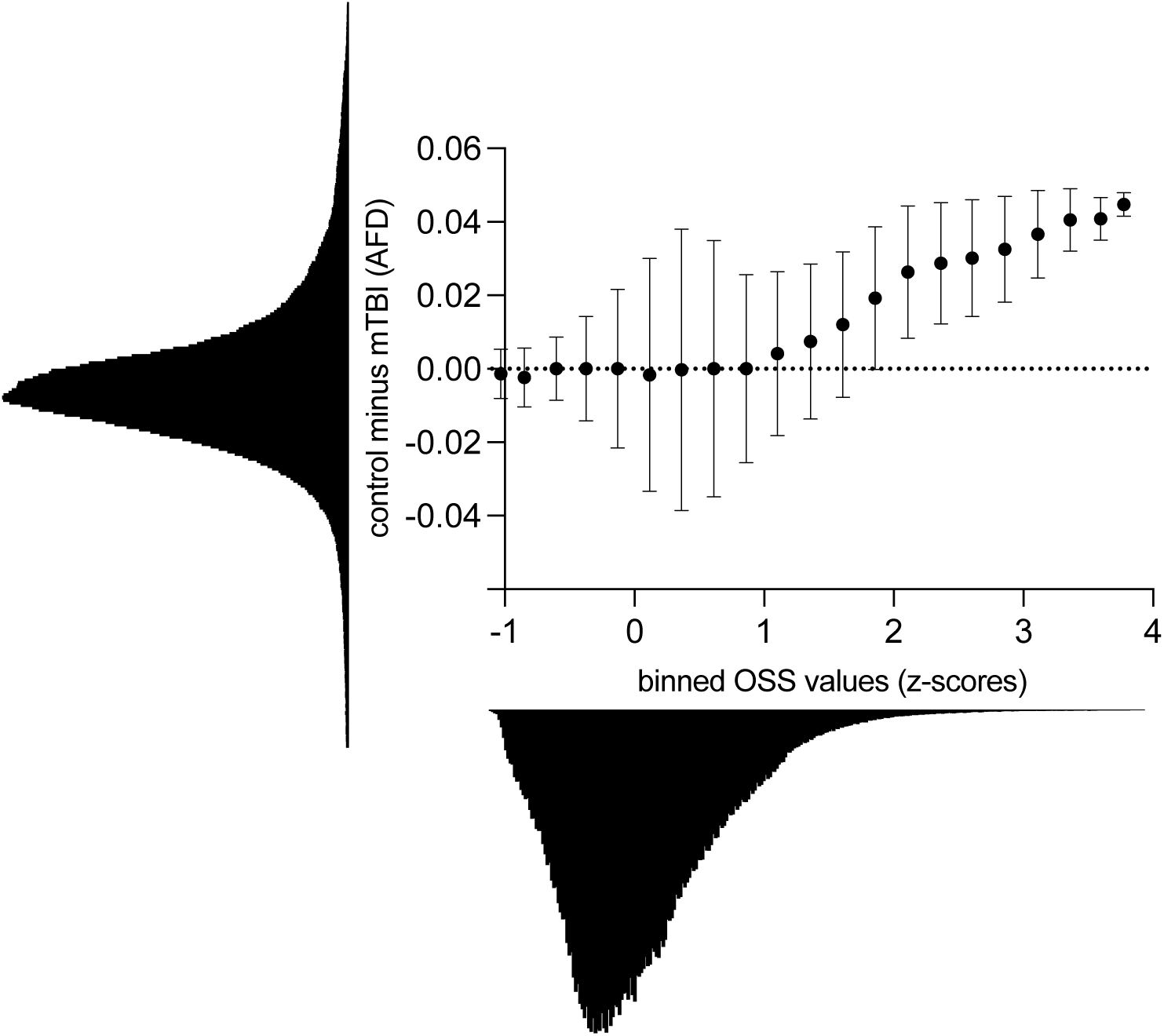
Threshold-agnostic whole-brain analyses demonstrate a positive relation between strain concentration and tissue integrity. Differences in AFD between mTBI and control participants (Control minus mTBI) are related to differences in shear strain (octahedral shear strain (OSS)) across the whole brain.

### Both differential energy arriving, and a differential reduction in dispersions of that energy, drive high strain concentration

As discussed in the introduction, high strain concentration in specific brain structures could be caused by two non-exclusive reasons: those structures experience high levels of shear wave forces (higher energy), and/or those regions are differentially stiff compared to surrounding brain structures, rendering them unable to dissipate the energy that arrives. We sought evidence for each of these alternatives in parallel, as they are non-exclusive biomechanical account of the differential injury onserved in high strain regions. We calculated shear strain energy 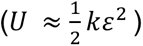 for those subjects (n=59) for which we had both stiffness (***κ***) and OSS (*ɛ*). We found that here is greater strain energy in high strain regions compared to the rest of the brain (t = 12.3, p < 0.0001; Wilcoxon matched-pairs rank test, p < 0.0001; Fig 4A). Those findings indicate that one cause of high strain is differential focusing of impact energy.

**Figure 4:**
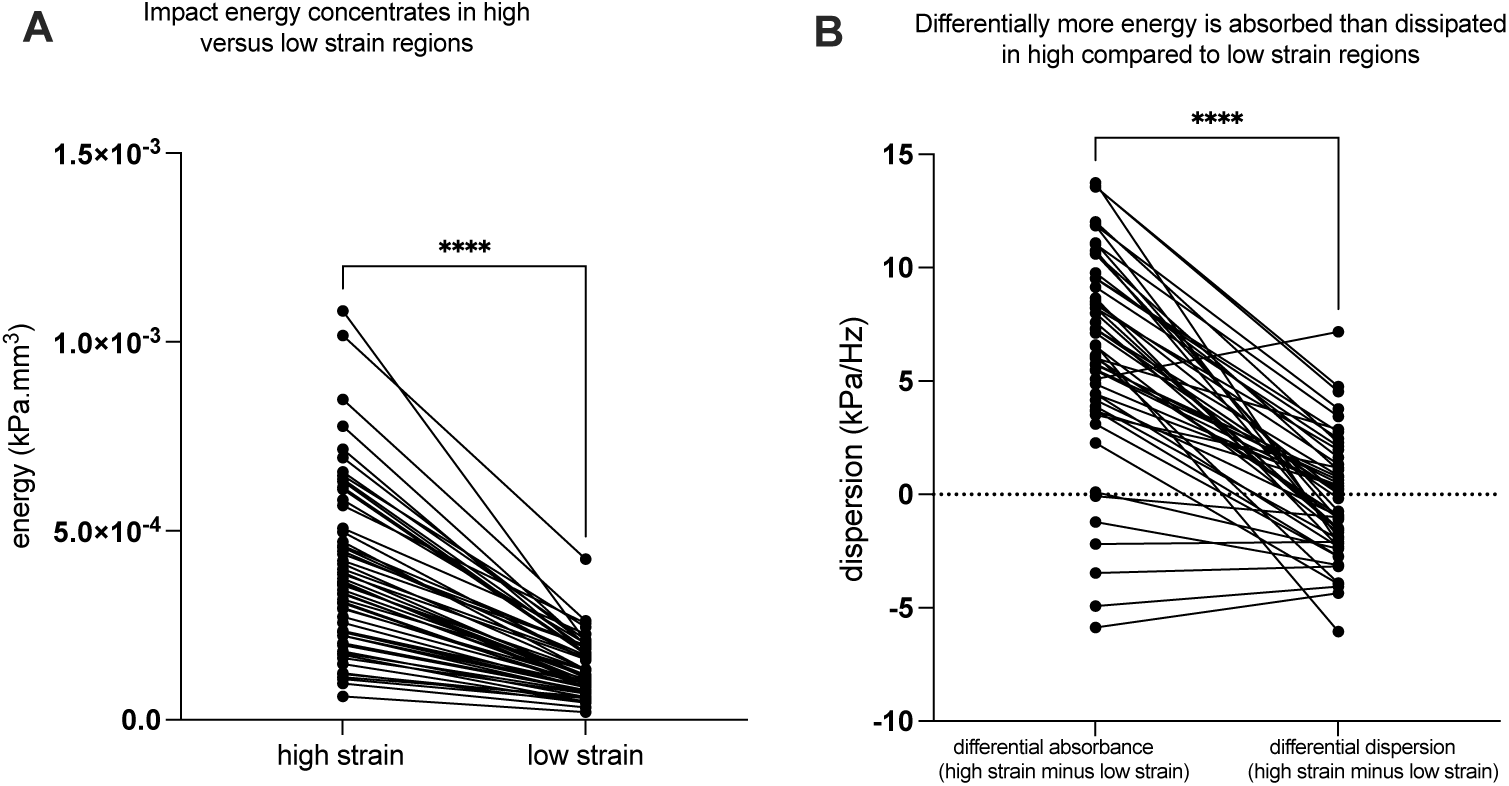
Differentially high strain is caused both by differential focusing of impact energy and reduced dissipation of that energy. ***A)*** Shear wave energy is differentially focused in regions of high strain concentration. ***B)*** The rate of energy absorbance is greater in regions with impact-location invariant high strain concentration compared to low strain regions, in nearly all of the participants. These findings mean that differentially high strain is due to *both* higher energy loading of those brain structures and the inability of those brain structures to dissipate that energy.

We also tested whether regions exhibiting differentially high strain (and high impact energy) also exhibit a reduced ability to dissipate the shear energy that arrives, compared to low-strain brain regions. To do that, we quantified the slopes of the storage and loss moduli as a function of frequency, taking advantage of multifrequency (30, 50, and 70Hz mechanical stimulation) Magnetic Resonance Elastography (MRE) data from a cohort of 54 participants. We found that high-strain brain regions absorb energy at a relatively greater rate than it is dissipated, compared to low strain regions (t = 12.6, p < 0.0001; Wilcoxon matched-pairs rank test, Figure 4B). This indicates that the viscoelastic properties of brain regions that exhibit injury in mTBI are differentially unable to dissipate energy from external skull loading.

## Discussion

We applied mechanics-based concepts to quantitatively test whether the patterns of force loading and force dissipation in the brain caused by mechanical skull stimulation predicts the locations of tissue disruption in mTBI. We used MRE to first show that the midbrain, cerebellum, mesial temporal lobe structures and sulcul depths exhibit differentially high strain concentration, or force loading (Fig 1A). Those regions are known for exhibiting tissue disruption in mTBI, and disruption to the normal functions of those regions explains the common signs and symptoms of mTBI ^11,12^. To empirically validate the biomechanical measurements we obtained in healthy participants using MRE, we tested if regions exhibiting high strain concentration are also those that exhibit reductions in tissue integrity in acute, sub-acute and chronic cohorts of mTBI participants (compared to respective control groups; Fig 2A and B). We observed a consistent pattern of differential injury in acute, sub-acute, and chronic mTBI patients compared to controls, in high-strain regions compared to low-strain regions. That relation between regions exhibiting high strain concentration (measured with MRE in healthy controls) and those showing differential tissue damage (measured with dMRI in mTBI participants) was present in whole-brain threshold-agnostic analyses in each of the post-concussive cohorts (acute, sub-acute, chronic) (Fig 3). Finally, we sought positive evidence for each of two non-exclusive biomechanical reasons for high strain concentration in specific brain structures: shear waves from skull impacts differentially concentrate in those regions, and/or, those regions are less able to dissipate the shear wave energy that arrives. We found evidence that there is indeed differential focusing of external shear forces leading to higher energy transmission to high-strain regions (Fig 4A), and that those regions absorb more energy than they dissipate, relative to low strain regions (Fig 4B). Overall, our findings ground a theoretical framework in a quantitative approach for testing how biomechanical principles drive the concentration of tissue disruption in specific brain regions associated with functions such as consciousness, balance, and memory in m/TBI. This *in vivo* approach in humans connects with a rich tradition of studying the histopathology of TBI-associated injury in post mortem human brains, and provides a biomechanical framework with which to understand the structural causes of the most common signs and symptoms associated with mTBI.

Previous studies have highlighted the biomechanical vulnerability of midline structures to injury in the setting of TBI. The brainstem, for example, is believed to be susceptible to TBI due to its location and geometry relative to the cortex, resulting in differential sensitivity to rotational loading ^13^. Finite Element Modelling (FEM) of video recordings of concussive episodes in the NFL have inferred rates of high strain in the midbrain, thalamus, and hippocampal/parahippocampal gyrus, accounting for concussions observed in American football players ^32^. Another study involving a comprehensive review of 1280 National Football League games identified 20 head impacts displaying evident signs of loss of consciousness, 21 with abnormal posturing, and 41 control impacts devoid of observable neurological signs for detailed video analysis. Those videos were used to guide physical reconstructions of the impacts in order to estimate impact kinematics ^11^. That detailed analysis revealed that impacts associated with loss of consciousness displayed significantly higher head acceleration and brain deformation in various regions, notably the midbrain and cerebellar areas, suggesting biomechanical vulnerability of brainstem nuclei that are instrumental in maintaining consciousness. In line with this, previous research from our group has shown that the amount of rotational loading experienced by collegiate American football players, as measured by helmet-worn accelerometers, predicts the extent of axonal injury in the midbrain, even in the absence of concussion^6^.

MRE allows for the quantitative measure of material and geometric discontinuities that causes shear strain in soft tissues such as the brain^33–36^. In line with established principles of mechanics, MRE-based measures in soft tissue indicate that strain concentrates interfaces between materials of different viscoelastic properties^37,38^. Our data align with prior unpublished transient MRE data obtained by the Mayo Clinic, showing strain concentration in the depths of sulci and the propagation of shear waves through midline structures in a healthy control subject ^39,40^. Although not explicitly stated by the authors, tagged-MRI data from the Bayly group also appears to align with our findings of strain concentration in the midbrain and within sulcal depths ^41–43^. Our work further aligns with modelling studies that show sulcal depths tend to experience relative high strains ^12^. Nevertheless, our present findings, to our knowledge, represent the first comprehensive demonstration that strain concentration maps, indexing biomechanical vulnerability, and obtained with *in vivo* measurements in humans, can identify regions that exhibit stereotyped injury in mild traumatic brain injury.

Mechanical theories predict that failure of a structure occurs when the deformation of that structure exceeds a threshold beyond which the material integrity of the structure is undermined. In biological organs, such as bones, such concentrations arise from stiffness discontinuities between heterogeneous structures ^44–46^. Our findings show that the brain, with its heterogeneous geometry, numerous interfaces, and viscoelastic discontinuities, has naturally occurring regions of differentially high strain concentration.

The existence of streotyped regions of high strain concentration in the human brain has implications for the interpretation of prior histopathology studies of brain injury caused by subclinical repetitive traumatic impacts, such as those sustained in the setting of contact sports and military operations ^23,31,47,48^. Brain regions representing differentially high levels of strain are likely to sustain differential disruption, regardless of impact magnitiude. Importantly, our findings also identify sulcal depths in addition to midline structures as sites of high strain concentration, reconciling prior finding of significant pathology in chronic traumatic encephalopathy, in both sulcal depths and midline structures ^49–54^. We also find that both gray and white matter are under disproportionate strain concentration, reconciling prior evidence for both gray and white matter changes in mTBI patients ^6,22,25,55–63^. In this regard, our findings underscore the importance of comprehensive assessment metrics that are sensitive to changes in both gray and white matter. In particular, the use of AFD is an important index that allows for *in vivo* quantification of both white and gray matter injury.

### Limitations

There are important limitations of our findings that motivate next steps. First, the magnitude of impact during mild Traumatic Brain Injury (mTBI) clearly exceeds the amplitude of the non-injurious actuation used during Magnetic Resonance Elastography (MRE) strain measurements. Replicating concussive loading magnitudes in living humans is unfeasible for obvious reasons. We have addressed this by using separate cohorts of mTBI patients to validate the premise that brain regions exhibiting differential strain concentration under conditions of non-injurious loading correspond to those observed in mTBI. It is also worth underlining that studies involving gyrencephalic animals and cadavers subjected to higher impact magnitudes support the observed differential strain concentration and pathology in the same regions we report ^9,13,15,26,64^. The correspondence of strain concentration across low and high magnitude controlled actuation of the human skull may be partially explained by the stereotyped response frequency of the brain, irrespective of input frequency and magnitude^64,65^. Future work should continue to explore how amplitude of skull actuation, always within the non-injurious range, modulates the pattern of findings we have reported.

A second limitation is that our study is limited by the absence of information regarding the direction of impact in our concussion cohort. We have addressed this by focusing on regions that exhibit strain concentration irrespective of the direction of impact. Nonetheless, our findeings compel a shift a in practice, which is to prospetively asses the direction of impact in clinical concussion assessment, in support of future tests of how strain concentration in specific structures is modulated by the direction of force propogation.

A final limitation concerns the heterogeneity of mTBI signs and symptoms. The framework we have articulated is largely based on the empirically established premise that there is a stereotyped pattern of injury in mTBI ^13–15,21,24^. At the same time, there is a wide range of inter-individual variability across the acute, sub-acute, and chronic phases of post-concussive recovery in terms of the pattern of signs and symptoms ^66–69^. Our findings reinforce what is common across m/TBI ^15,19,21^, in a way that is grounded in the biomechanics of shear wave propagation through tissue. The flip side of this is the exciting prospect that person-specific patterns of strain concentration may be inferred from individual variability in brain anatomy. That approach could move the field closer to developing biomechanically grounded prognostic models about where injury is predicted to differentially accumulate for a given individual, for a given head impact. By being able to identify the signal that is common and shared across variability in brain anatomy and skull-impact location, future work can isolate individual-specific variance in signs and symptoms and relate that to person-specific viscoelastic properties of individual brain anatomy, and clinical signs and symptoms.

## Significance Statements

- Magnetic Resonance Elastography uses mechanical actuation of the skull to measure how shear waves propograte through tissue *in vivo*
- Strain concentration is a quantitative biomechanical index that captures how energy is focused on specific brain structures after force-loading on the skull
- Brain regions with high strain concentration are sites of differential tissue injury in acute, sub-acute, and chronic mTBI
- In vivo analysis of strain concentration grounds a new biomechanical framework for understanding stereotyped and person-specific patterns of tissue injury in mTBI and TBI

## Authors’ Note and Disclosure

Preparation of this ms was supported, in part, by NIH Grants R01NS089069 and R01EY02853 to BZM. The authors are grateful to Dr. Jeff Bazarian and Kian Merchant for their assistance supporting IRB oversite for the healthy control diffusion MR data used to reference to mTBI diffusion MR data. AAH and BZM hold IP related to predicting injury from mTBI using MRI (US-20220039732-A1).

## Data materials and availability

All data needed to evaluate the conclusions in the paper are present in the paper and/or the Supplementary Materials.

---

**Supplemental Figure 1:**
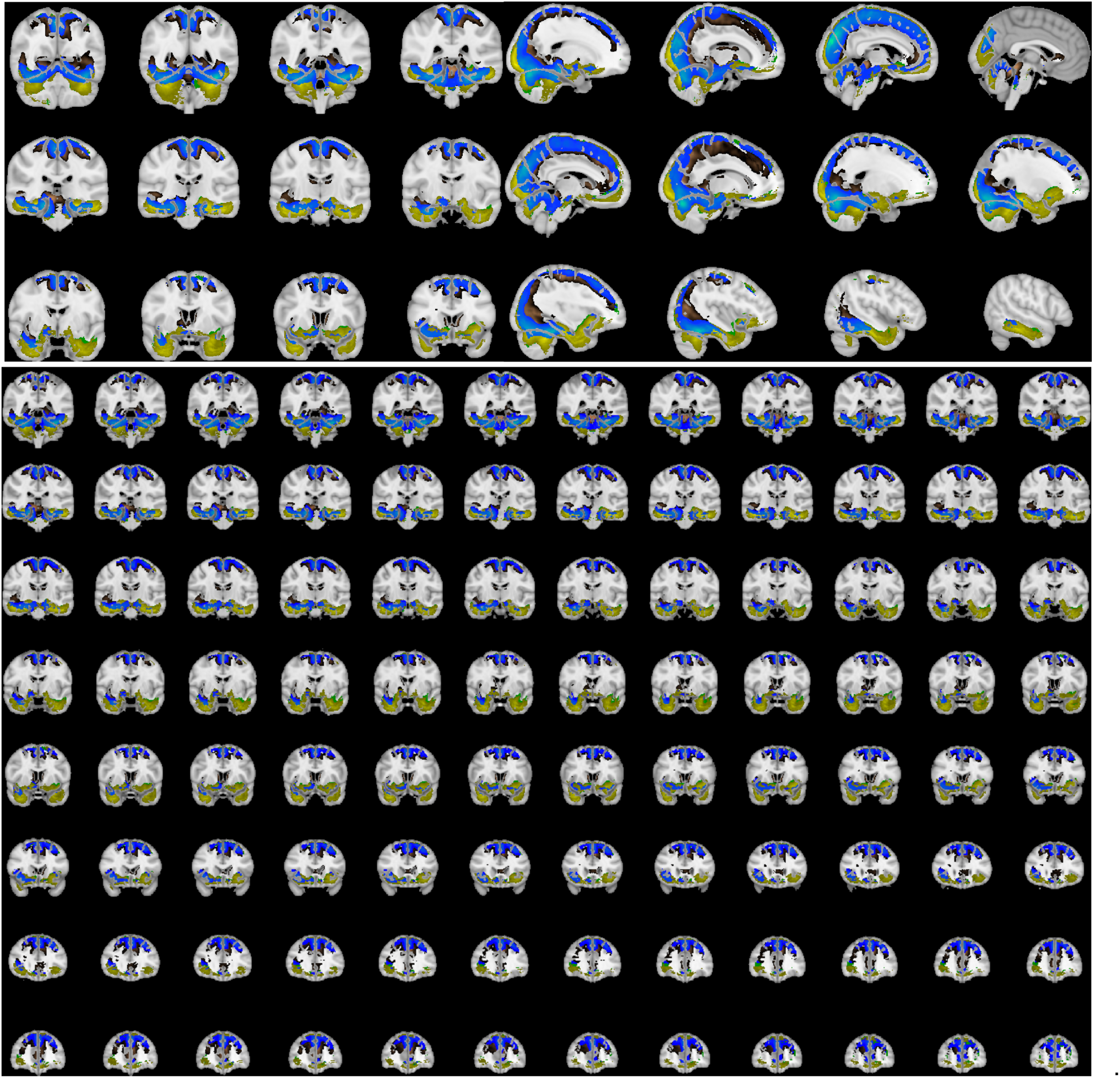
Overlap and dissociation of high strain regions for 30, 50 and 70 Hz occipital actuation MRE data. Strain concentration defined at a threshold of T = 4 (FDR-corrected q < .05) for 30 Hz (copper), 50 Hz (green), and 70 Hz (yellow) and intersection (blue). Blue colored regions experience differentially high strain concentration across all three tested frequencies.

